# Hepatocyte nuclear factor 4-mediated lipotoxicity provokes mitochondrial damage in peroxisome-deficient *pex19* mutants

**DOI:** 10.1101/071787

**Authors:** Margret H. Bülow, Julia Sellin, Christian Wingen, Deniz Senyilmaz, Dominic Gosejacob, Aurelio A. Teleman, Michael Hoch

## Abstract

Peroxisomes are important metabolic organelles involved in the catabolism of several lipid classes, e.g. very-long-chain fatty acids. Malfunction or absence of peroxisomes leads to accumulation of educts for peroxisomal β-oxidation and mitochondrial damage, resulting in fatal perturbation of metabolism. The impact of peroxisome deficiency on mitochondria is not elucidated yet. Here we present a model of Hepatocyte nuclear factor 4 (Hnf4)-induced lipotoxicity and accumulation of non-esterified fatty acids (NEFA) as the cause for mitochondrial damage in consequence of peroxisome loss in a Peroxin19 (*pex19*) mutant. Hyperactive Hnf4 signaling leads to upregulation of *lipase 3* and enzymes for mitochondrial β-oxidation. This results in enhanced lipolysis, elevated concentrations of NEFA, maximal β-oxidation and mitochondrial swelling. NEFA are ligands for Hnf4 and further enhance its activity. By genetic removal of Hnf4 in *pex19* mutants, lipotoxicity and mitochondrial swelling are reduced and their survival is rescued.

**Author summary:** Peroxisomes are cell organelles which play a major role in lipid metabolism. They interact with mitochondria, the organelles which are responsible for cellular energy production. Loss of peroxisomes, as it occurs in the rare, inheritable human disease class of Peroxisome Biogenesis Disorders, is lethal. Over the past couple of years, a number of studies showed that peroxisome loss leads to mitochondrial damage as a secondary consequence, but the underlying mechanism has not been understood yet. In our study, we use a mutant of the fruitfly *Drosophila melanogaster* as a model for Peroxisome Biogenesis Disorders and find that a protein called Hepatocyte nuclear factor 4 is hyperactive upon peroxisome loss, which provokes the mobilization of storage fat and, as a consequence, the accumulation of toxic free fatty acids. These enter the mitochondria, but cannot be used for energy gain. Free fatty acids are then trapped in the mitochondria and lead to their swelling and damage, which provides an explanation for mitochondrial defects in Peroxisomal Biogenesis Disorders. Genetic reduction of Hepatocyte nuclear factor 4 activity rescues the viability of the peroxisome mutant by reducing the accumulation of free fatty acids and the subsequent mitochondrial damage, which might provide a novel target for therapy development.

## Abbreviations

4E-bp: 4E-binding protein
Acsl: Acyl-CoA synthetase long-chain
Bmm: Brummer lipase
CPT-1: Carnitine-palmitoyl-transferase 1
FA: Fatty acid
FAME: Fatty acid methyl ester
Fas: Fatty acid synthase
FITC: Fluorescein isothiocyanate
HexC: Hexokinase C
Hnf4: Hepatocyte nuclear factor 4
InR: Insulin receptor
LCFA: Long chain fatty acid
Lip3: Lipase 3
MCFA: Medium chain fatty acid
NEFA: Non-esterified fatty acid
PBD: Peroxisomal biogenesis disorder
Pepck: Phosphoenolpyruvate carboxykinase
Pex: Peroxin
Pex19: Peroxin 19
PMP: Peroxisomal membrane protein
TMRE: Tetramethylrhodamine, ethyl ester
VLCFA: Very long chain fatty acid
Yip2: Yippee interacting protein 2

## Introduction

Peroxisomes, while rather simply structured organelles delimited by a single membrane, harbor complex metabolic functions, which are still incompletely understood. In mammalian cells, they are involved in the β-oxidation of very long chain fatty acids (VLCFA), the formation of ether phospholipids (like plasmalogens), the catabolism of branched chain fatty acids, the production of bile acids, polyamine oxidation and amino acid catabolism. Furthermore, they exhibit a functional interplay with mitochondria by employing both shared and coordinated metabolic pathways [1]-[3]. They interact via mitochondria derived vesicles (MDVs), which allow for the exchange of lipid as well as protein content between the two compartments [4], and they share regulators controlling organelle biogenesis [5] and fission [6], [7]. VLCFA and branched-chain FA have to undergo β-oxidation in peroxisomes before they can be metabolized in the mitochondria [8]. The interplay between peroxisomes and mitochondria is illustrated by the fact that loss of peroxisomes leads to defects in mitochondrial metabolism and structure [9]-[11]. The mechanism behind these defects, however, is not understood.

Peroxisomes primarily regulate their number by growth and division of preexisting organelles, similar to mitochondria, or by de novo biogenesis via pre-peroxisomal vesicles through budding off from the endoplasmic reticulum (ER) [12]-[15]. The machinery involved in the inheritance, assembly, division and maintenance of peroxisomes is encoded by *peroxin (pex*) genes, which are highly conserved in evolution from yeast to mammals [16]. Pex19 is a predominantly cytoplasmic peroxisomal core factor and essential for both the import of peroxisomal membrane proteins (PMPs) and the de novo formation of peroxisomes [17], [18]. Together with Pex3 and Pex16, it is responsible for the translocation of membrane proteins and membrane vesicle assembly [19]. Mutations in *pex* genes lead to the loss of peroxisomes and cause peroxisome biogenesis disorders (PBDs), and *pex19* loss of function specifically leads to Zellweger Syndrome, the severest form of PBDs. Recently, Pex19 has been found to be required not only for peroxisome biogenesis, but also lipid droplet formation from the ER [20]. The peroxisomal biogenesis and assembly machinery as well as metabolic function are well conserved in *Drosophila melanogaster* [21]-[24].

Hepatocyte nuclear factor 4 alpha (HNFα) is a ligand-regulated transcription factor which acts as a regulator of lipid metabolism in the mammalian liver [25]. It is conserved in invertebrates, and in *D. melanogaster*, the single orthologue Hnf4 acts as an important lipid sensor. Hnf4 is activated by binding of non-esterified fatty acids (NEFA), which are generated by lipolysis of storage fat upon starvation. Binding of NEFA leads to nuclear translocation of Hnf4 and the transcription of its target genes, which further triggers lipolysis and mitochondrial β-oxidation [26].

Here we present our study on a *D. melanogaster pex19* mutant, which recapitulates all major hallmarks of Zellweger Syndrome, such as absence of peroxisomes and consequently VLCFA accumulation, mitochondrial defects, neurodegeneration, and early lethality, therefore serving as a model for PBDs. This simple and genetically tractable model system was used in order to identify pathological factors contributing to mitochondrial dysfunction. We were able to identify one major pathological cascade as a result of peroxisome loss: Hyperactive Hnf4 signaling and severely increased lipolysis in *pex19* mutants promote mitochondrial damage by increasing mitotoxic NEFA levels. Our data therefore contribute to the current efforts in the field.

## Results

### Neurodegeneration and aberrant lipid metabolism in *pex19* mutants

The single Pex19 orthologue in *D. melanogaster* is well conserved on the sequence and structural level. It contains the typical Pex19 core domain [27] (Fig. 1a) and a highly conserved Pex3 binding domain, both required for its activity as cytoplasmic shuttle receptor of the peroxisome import machinery. We generated a deletion of the coding region of the gene by imprecise excision (Fig. 1b), and molecularly characterized and identified the resulting *pex19*^ΔF7^ flies as null mutants (subsequently referred to as *pex19* mutants). Maternal-zygotic *pex19* mutants die as embryos, and ultrastructural analysis revealed the absence of peroxisomes (Fig. 1c 1-2). In zygotic mutants, peroxisomes get lost during larval development in these animals (Fig. 1c 3-4), with impact on their survival: while most larvae develop to pupae (~70%), only 20% adults develop and most of them die during the first 24 hours (Fig. 1d). The few adult escapers (“surviving adults”, Fig. 1d) have high numbers of apoptotic cells in their brains (Fig. 1c 5-6).

**Figure 1:**
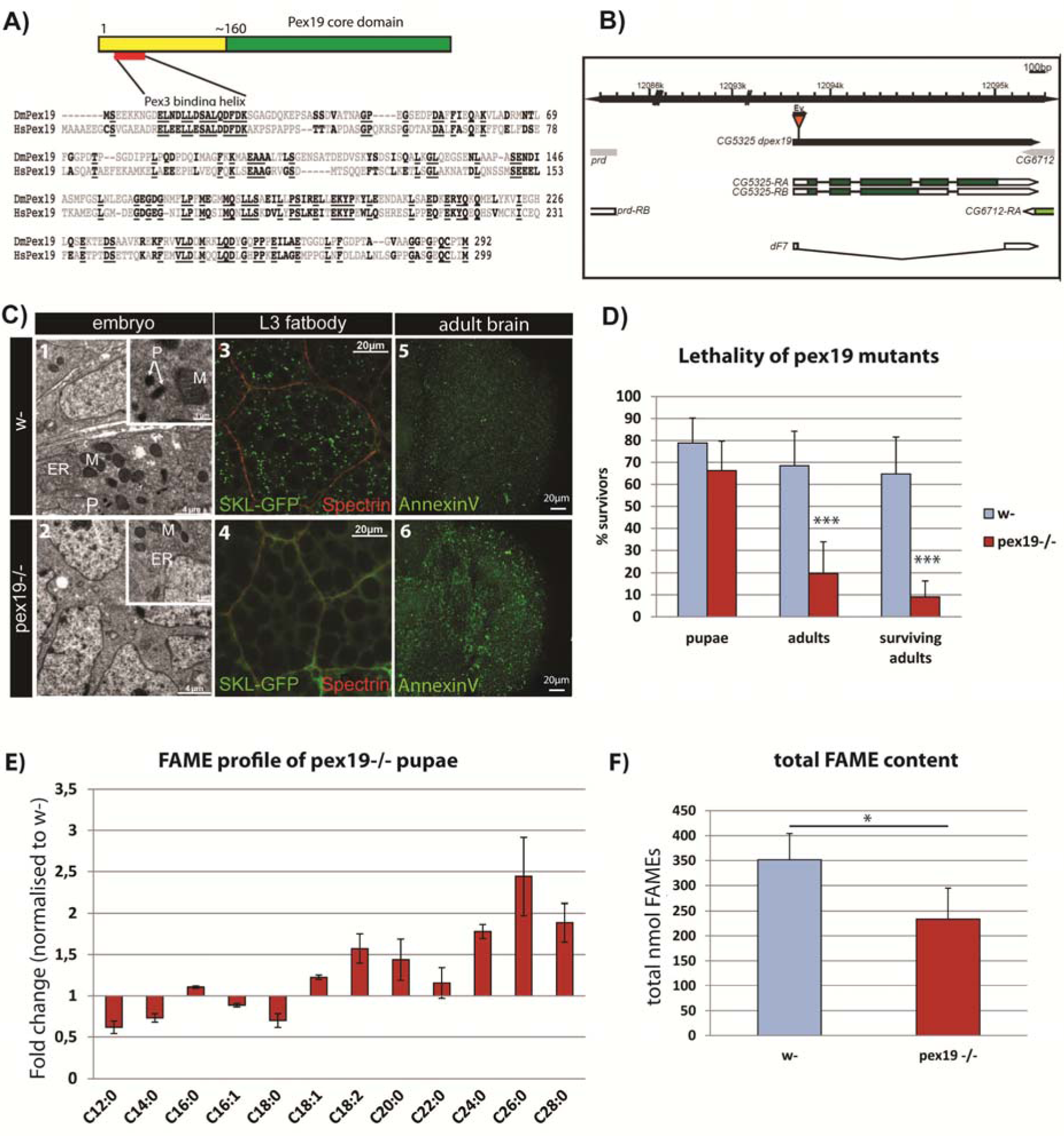
**A)**Protein structure and sequence of Pex19 from human (HsPex19) and *Drosophila melanogaster* (DmPex19) and their conservation. **B)** Schematic representation of the *pex19* gene locus and the deletion in the *pex19*^ΔF7^ mutant. **C)** Peroxisomes are absent from *pex19*-/- animals. Ultrastructural analysis of maternal-zygotic *pex19*-/- mutant embryos (2, compare to wildtype in 1) shows loss of peroxisomes. P: peroxisomes, M: mitochondria, ER: endoplasmatic reticulum. Zygotic pex19-/- larvae do not have peroxisomes either, as shown by crossing in the peroxisome marker SKL-GFP: Loss of GFP positive punctae in 4 indicates loss of peroxisomes (compare to wildtype in 3). AnnexinV-FITC staining of adult optic lobes indicate neurodegeneration by high numbers of apoptotic cells in pex19 mutant brains (6, compare to 5). **D)** Lethality profile of *pex19-/-* mutants, indicating the number of pupae, adults including pharates and viable adults which survived for more than 24 hours. **E)** Fatty acid methyl esters (FAME) from *pex19*-/- pupae, normalized to w- control. **F)** Sum of lipids measured in the FAME profile. Scale bars as indicated. Error bars represent standard deviation (SD). *p<0,05, ***p<0,001 (Student’s t-test)

A genetic rescue with a Pex19 expression construct could be achieved with the ubiquitous tubulin-Gal4 driver and with pumpless-Gal4, which drives expression in the fatbody and gut. Expression with neuronal (elav-Gal4) or glia (repo-Gal4) drivers failed to rescue the lethality of *pex19* mutants, which highlights an important role for Pex19 in metabolic organs rather than the CNS (supplemental Fig. S1a).

Consistent with a loss of peroxisomal function, GC/MS analysis of fatty acid methylesters (FAMEs) prepared from pupae shows that lipids containing VLCFA are present at increased levels in flies lacking peroxisomes. In addition, we found that lipids containing MCFA and LCFA with C12 – C18 chain lengths are reduced (Fig. 1e). Also, the total amount of FAMEs is lowered (Fig. 1f). VLCFA are present at ~ 0,1 - 0,5 nmol per animal and their contribution to energy gain is therefore negligible, while M- and LCFA are present at ~2 - 125 nmol per animal (supplemental Fig. S1b) and provide fuel for energy gain by the mitochondria. The shortage in MCFA and the reduction in total lipids is accompanied by a low energy status: AMP kinase, a sensor for the AMP/ATP ratio, is phosphorylated and therefore indicates low ATP levels [28] (supplemental Fig. S1c).

### Gene expression of metabolic enzymes is altered in pex19 mutants

To further characterise the lipid imbalance, we analyzed the expression of several genes encoding for metabolic enzymes reportedly regulated on the transcript level [29], as well as other enzymes of interest (Table 1). We suspected that *pex19* mutants are in a state of starvation (low ATP levels and lack of MCFA), but instead found that insulin signaling is elevated, as indicated by reduced expression of dFoxo target genes (*4E-BP* and Insulin Receptor, *InR*). When analyzing the transcript levels of lipases, we found most of them reduced, with the exception of *lipase 3,* which was strongly elevated (~250 fold compared to w- control). Unlike *lipase 4* [30] and the ATGL homolog *brumer* [31], *lip3* is not a dFoxo target gene, although it is upregulated in starved animals [32]. High expression of *lip3* is therefore indicating a starvation situation in *pex19* mutants in spite of the fact that insulin signaling is high. This surprising contradiction raises the question if lipid sensing is impaired in *pex19* mutants, since lipolysis is upregulated in spite of high insulin signaling.

**Table 1:**
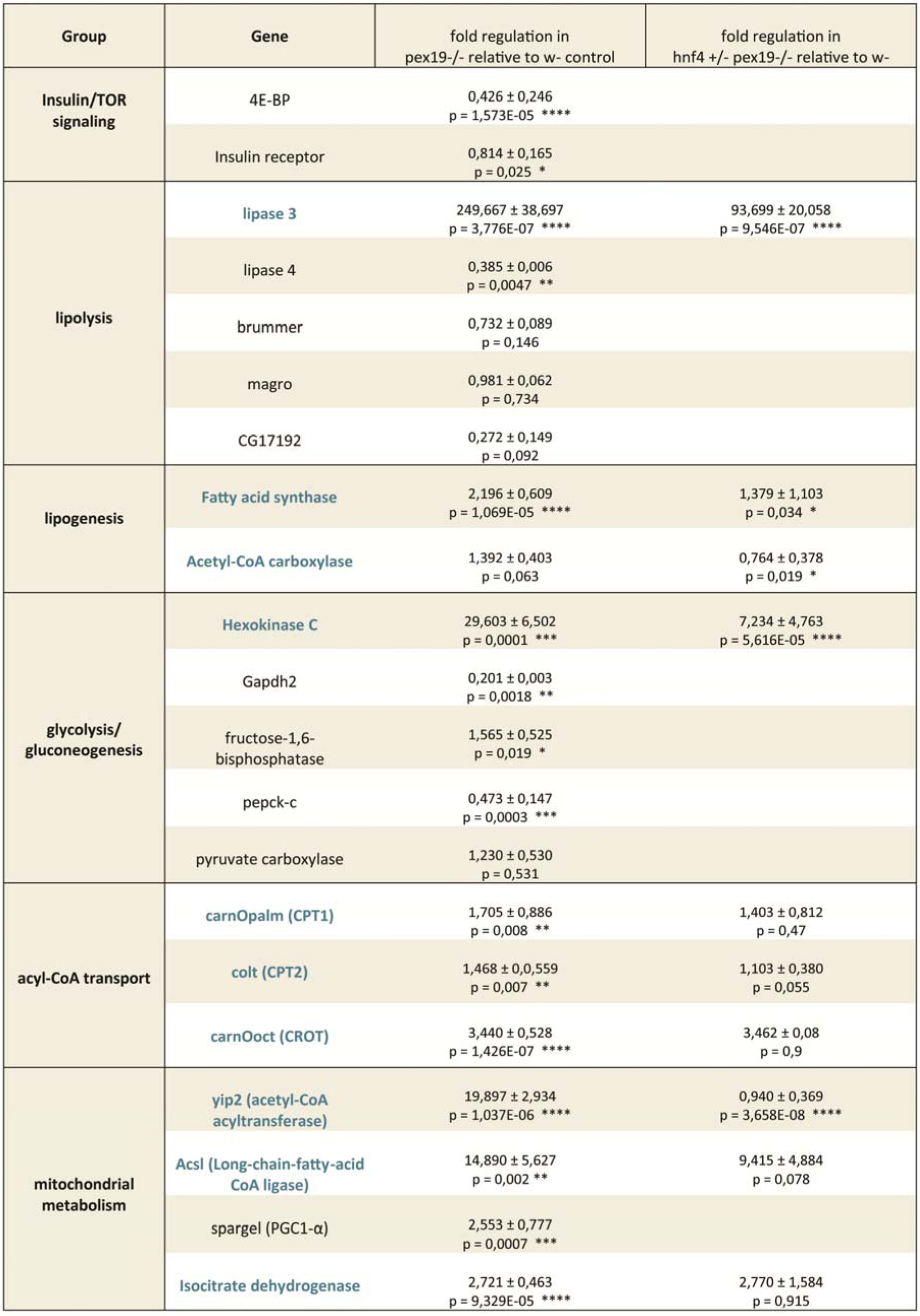
Real-time qPCR analysis of genes encoding for metabolic enzymes. Δ_cq_ values are normalized to w- (ΔΔC_q_or fold regulation). Putative Hnf4 target genes are typed in blue. CPTI: carnitine palmitoyltransferase, CROT: carnitine octanoyltransferase, PGC1-α: Peroxisome proliferator-activated receptor gamma coactivator 1-alpha. Significance for *pex19-/-* values tested against w- values and for *hnf4+/- pex19-/-* values against *pex19-/-.* ± indicates SD. *p<0,05, **p<0,01, ***p<0,001, ****p<0,0001

Among the glycolytic and gluconeogenetic enzymes, we found *hexokinase c*, which catalyzes the initial step of glycolysis (conversion of glucose to glucose-6-phosphate), ~30 fold upregulated. We analyzed 3 genes involved in mitochondrial acyl-CoA import, carnitine-O-octanoyltransferase (CROT) and the carnitine-palmitoyltransferases 1 and 2, and found all of them upregulated in *pex19* mutants. Other genes for enzymes involved in mitochondrial metabolism (β-oxidation, TCA cycle), most prominently the acetyl-CoA acyltransferase yip2 (yippee interacting protein 2, ~20 fold) and the long-chain fatty acid CoA ligase (Acsl, ~15 fold), were also upregulated. Strikingly, most of the upregulated genes in *pex19* mutants are targets of the Hepatocyte nuclear factor 4 (Hnf4), a transcription factor which regulates metabolic processes such as lipolysis and mitochondrial fatty acid β-oxidation in response to starvation [26], and which stimulates insulin secretion [33]. This prompted us to investigate Hnf4 in the context of peroxisome deficiency.

### Hnf4 is hyperactive in *pex19* mutants and responsible for their reduced viability

We used a LacZ reporter under the control of a heat-shock inducible Hnf4:Gal4 fusion protein [26], which did not show Hnf4 activity in fatbodies of 3rd instar wild type larvae. By contrast, Hnf4 activity was observed in *pex19* mutants (Fig. 2a). We reanalyzed the Hnf4 target genes among the metabolic enzymes displayed in Table 1 and found that *lip3*, *fas*, *acc*, *hexC* and *yip2* are indeed significantly downregulated in *hnf4, pex19* mutants in comparison to *pex19* mutants (Fig. 2b). We concluded that counteracting Hnf4 hyperactivity by introducing a heterozygous mutation in *pex19* mutants normalizes the high expression of important Hnf4-regulated metabolic enzymes.

**Figure 2:**
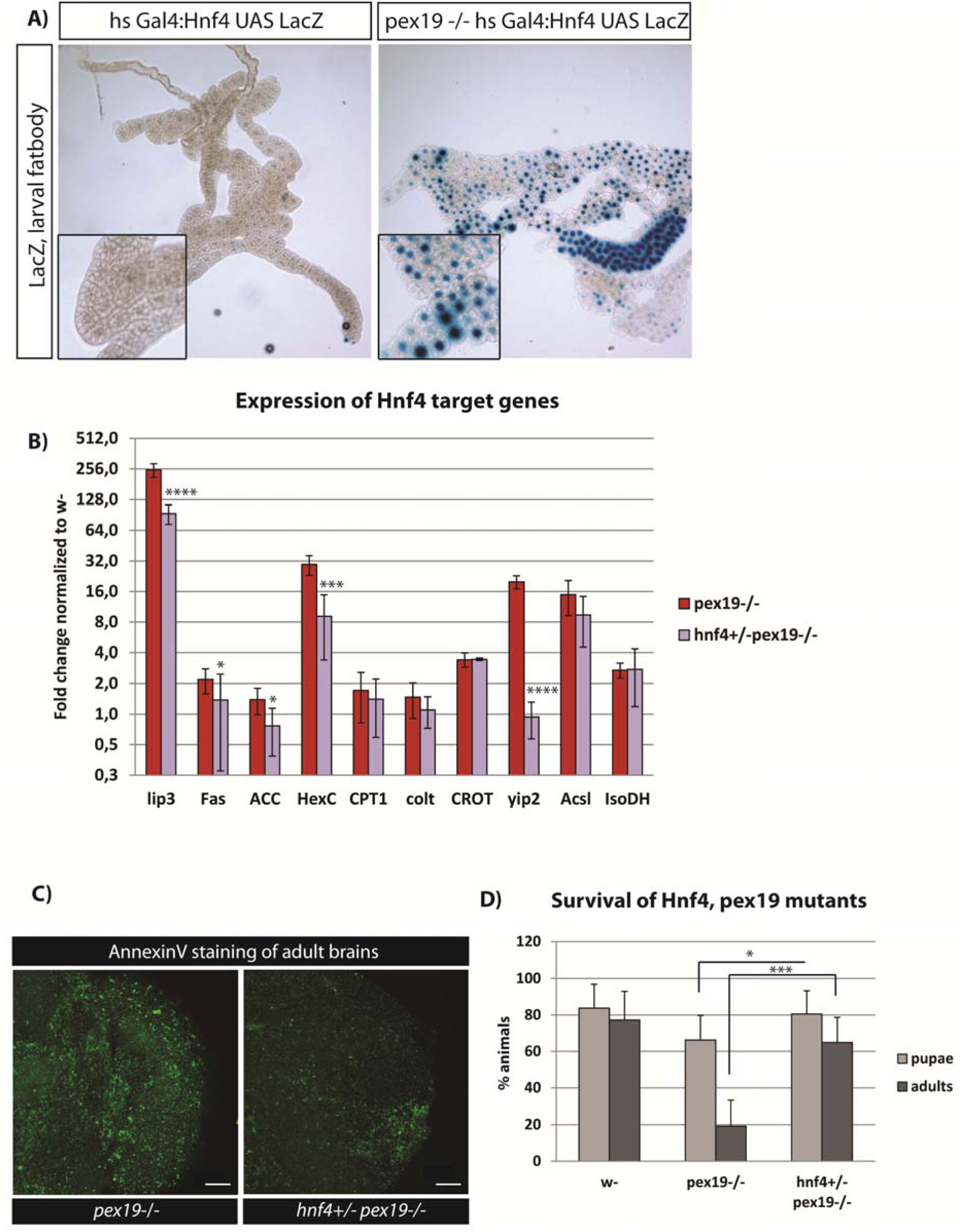
**A)** An HNF4 reporter, inducible by heat-shock, which expresses LacZ, was crossed into the *pex19* mutant background. HNF4 is induced in fatbodies of 3rd instar larvae in *pex19* mutants. **B)** Representation of the putative Hnf4 target genes: all of the measured Hnf4 target genes are upregulated in *pex19* mutants and most are downregulated in *hnf4+/- pex19-/-.* **C)** AnnexinV-FITC staining of adult optic lobes. Scale bars represent 20 µm. **D)** Lethality profile of *hnf4+/- pex19-/-* mutants, indicating the number of pupae, adults including pharates, and viable adults which survived for more than 24 hours. Error bars indicate SD. *p<0,05, ***p<0,001, ****p<0,0001

Furthermore, we assessed the neurodegeneration of *hnf4, pex19* mutants by staining brains of 5 day old adults with AnnexinV-FITC and found that the number of apoptotic cells decreases in comparison to *pex19* mutants (Fig. 2c).

Next, we tested the lethality of *hnf4, pex19* mutants and found that *pex19* mutants can be rescued to adulthood by removal of one copy of Hnf4. The resulting *hnf4, pex19* mutants showed a rate of hatched adults of >60% (Fig. 2d). Adult *hnf4, pex19* mutants were viable and fertile, but died 2-3 weeks after hatching from the pupa. This result indicates that elevated Hnf4 signaling has a major contribution to the lethality of *pex19* mutants.

### Hnf4 induces lipolysis and β-oxidation in *pex19* mutants

Hnf4 is an important lipid sensor in *D. melanogaster*. It acts as a receptor for non-esterified fatty acids (NEFA) and, in consequence of NEFA-binding, regulates the transcription of lipolytic and β-oxidation enzymes [26]. Intriguingly, the lipolytic enzyme *lipase 3* is extremely highly upregulated in *pex19* mutants, which prompted us to investigate the gut lipid stores. We stained neutral lipids with OilRed O in gut tissue of 3rd instar larvae and found that gut lipid stores are empty in *pex19* mutants while their oenocytes are filled with lipid droplets (supplemental Fig. S1d), which indicates lipid mobilization [34], consistent with lowered amounts of M- and LCFA and overall FAME content. Gut lipid filling is restored in *hnf4, pex19* double mutants, indicating that the lipolytic phenotype of *pex19* mutants is indeed connected to Hnf4 (Fig 3a).

**Figure 3:**
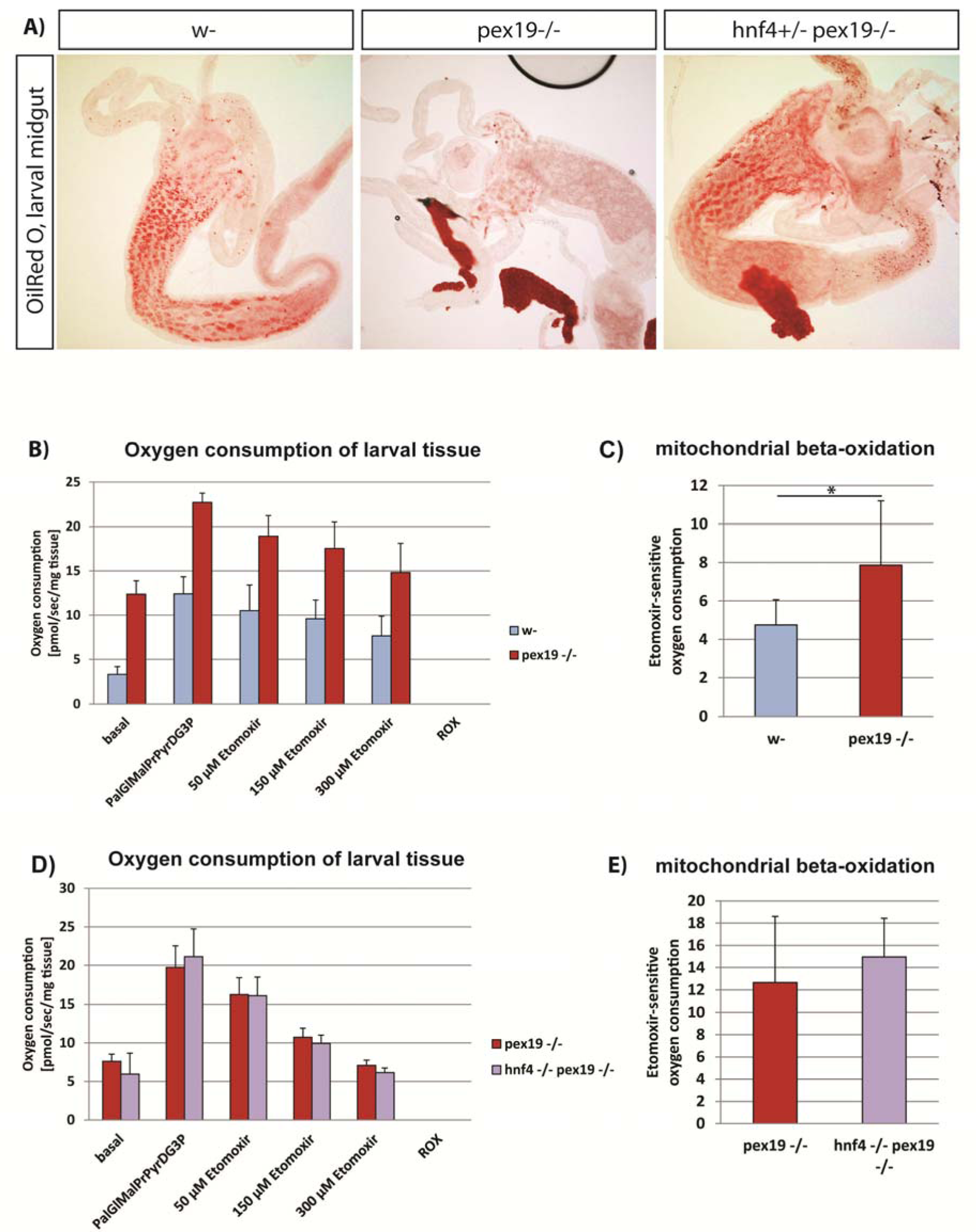
**A)** OilRed O staining of neutral lipids in anterior midguts of 3rd instar larvae. **B)** Oxygen consumption levels of wildtype and *pex19-/-* larval tissue (Pal: palmitate, Gl:glutamate, Mal:malate, Pr:proline, Pyr:pyruvate, D:ADP, G3P:glycerol-3-phosphate). **C)** Etomoxir-sensitive mitochondrial β-oxidation rate. **D)** Oxygen consumption levels of *pex19-/-* and *hnf4-/- pex19-/-* larvae. **E)** mitochondrial β- oxidation rate. Error bars represent SD. *p<0,05

Next, we analyzed mitochondrial β-oxidation in permeabilized third instar larvae using a Clark electrode (Oxygraph). β-oxidation levels were directly quantified as the amount of oxygen consumption (in the presence of palmitoyl-CoA substrate, the necessary TCA intermediates, and ADP) that is sensitive to etomoxir, a standard carnitine palmitoyl transferase (CPT-I) inhibitor [35], which has been used successfully in *D. melanogaster* before [36], [37]. We found that *pex19* mutants have elevated levels of etomoxir-sensitive oxygen consumption as a measure for mitochondrial β-oxidation, which is in line with high Hnf4 activity, increased lipolysis and reduction of M- and LCFA (Fig. 3b, c). To exclude that this is due to an increase of mitochondrial abundance, we measured mtDNA and citrate synthase activity, which are similar in *pex19* mutant and wildtype larvae (supplemental Fig. S2a, b), indicating that the increased oxygen consumption is due to increased mitochondrial flux, and not increased amounts of mitochondria. We measured the oxygen consumption in *hnf4, pex19* double mutants and found that removal of Hnf4 leaves mitochondrial β-oxidation at high levels (Fig. 3d, e). Since the expression of genes relevant for β-oxidation is lowered in *hnf4, pex19* compared to *pex19* mutants, this result suggests that the β-oxidation machinery is at maximal capacity.

### Hnf4-induced lipase 3 leads to mitochondrial swelling and high amounts of NEFA

We hypothesized that high Hnf4 signaling in *pex19* mutants leads to enhanced *lipase 3* activity and lipolysis, resulting in too high levels of free fatty acids (non-esterified fatty acids, NEFA) and maximum mitochondrial β-oxidation as a compensatory mechanism. When too many fatty acids are set free from the lipid stores by lipases, they can enter the mitochondrion without being activated with CoA, which prevents their β-oxidation and leads to mitochondrial swelling and damage. To analyze the NEFA content of tissue of *pex19* mutant larvae, we developed an adapted protocol of the copper-triethanolamine method [38] suitable for *D. melanogaster* tissue. We found that the NEFA content is about twice as high in *pex19* samples compared to w-, while *hnf4, pex19* mutants have wildtypic levels of NEFA (Fig. 4a).

**Figure 4:**
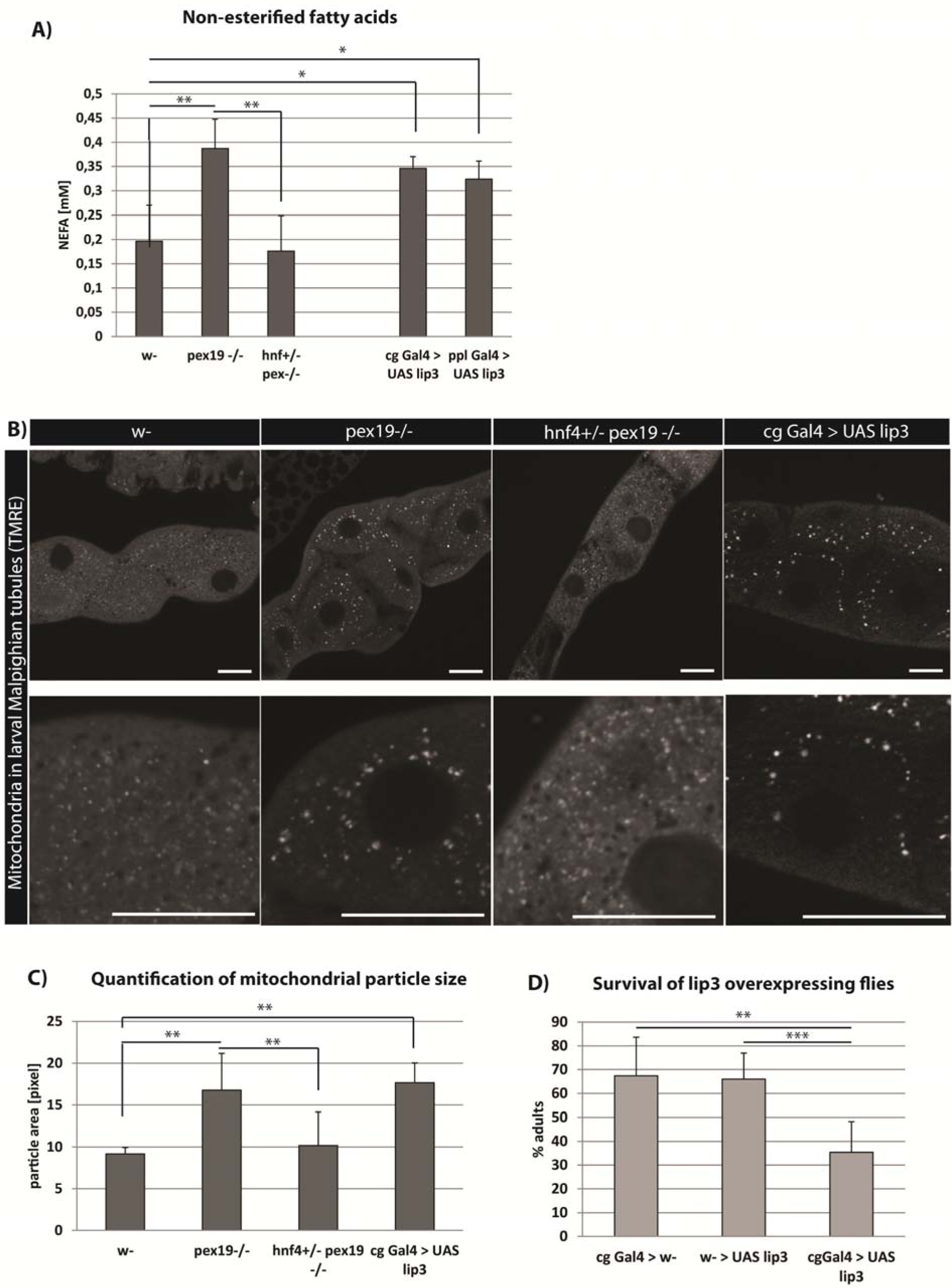
**A)** Amount of non-esterified fatty acids (NEFA) in whole 3rd instar larvae. **B)** TMRE staining to visualize active mitochondria in Malpighian tubules of 3rd instar larvae. Scale bars represent 20 µm. **C)** Quantification of the average particle area of TMRE-positive compartment. **D)**Flies overexpressing lipase 3 in the fatbody show reduced survival in comparison to controls. Error bars represent SD. *p<0,05, **p<0,01

To test whether NEFA accumulation was due to high *lipase 3* expression, we measured the NEFA levels in larvae overexpressing *lipase 3* under the control of the fatbody drivers pumpless (ppl)-Gal4 or combgap (cg)-Gal4. These animals show high expression of *lipase 3* relative to wildtype levels, comparable to *pex19* mutants (100-150fold, supplemental Fig. S2c). With both drivers, *lipase 3* overexpression leads to elevated NEFA levels comparable to those in *pex19* mutants (Fig. 4A).

To further strengthen the hypothesis that NEFA accumulation in *pex19* mutants leads to impaired mitochondrial function, we stained mitochondria with tetramethylrhodamine ethylester (TMRE), a dye that is readily incorporated by active mitochondria and indicative of their membrane potential. In Malpighian tubules of 3^rd^ instar larvae, mitochondria were reduced in number and enlarged in size in *pex19* mutants. This mitochondrial swelling was ameliorated in *hnf4,pex19* mutants (Fig. 4B, C). To test whether mitochondrial swelling was due to high NEFA levels in response to elevated *lipase 3* expression, we stained tissue of larvae expressing *lip3* under the control of the cg-Gal4 fatbody driver, and found that mitochondrial swelling occurs also in these animals. Furthermore, they show significantly reduced viability compared to control flies (Fig. 4D), although not to the same extent as *pex19* mutants.

Taken together, our data show that elevated Hnf4 signaling leads to high expression levels of *lipase 3* and enhanced lipolysis, resulting in accumulation of NEFA, which enter the mitochondria where they lead to mitochondrial swelling and dysfunction. A MitoSOX Red staining confirmed high ROS production by mitochondria in *pex19* mutants (supplemental Fig. S2E).NEFA act as ligands for Hnf4 and thus activate it further, thereby causing a self-sustained downward spiral culminating in energy deficit due to mitochondrial damage and ultimately death of the organism.

### NEFA accumulation in response to peroxisome loss is conserved between flies and humans

In order to assess the cross-species portability of our findings, we analyzed the mitochondrial morphology and NEFA content in a human fibroblast line from a Zellweger Syndrome patient with a mutation in pex19 [39] (Δ19T). While mitochondria appear small and network-like in a control cell line, they were fragmented and swollen in the Δ19T cells (Fig. 5A). Furthermore, NEFA were increased in these cells in comparison to the control cell line (Fig. 5B). Our results suggest a similar mechanism of the impact of *pex19* mutation on mitochondria and the pathology of the disease as observed in our *D. melanogaster* mutant.

**Figure 5:**
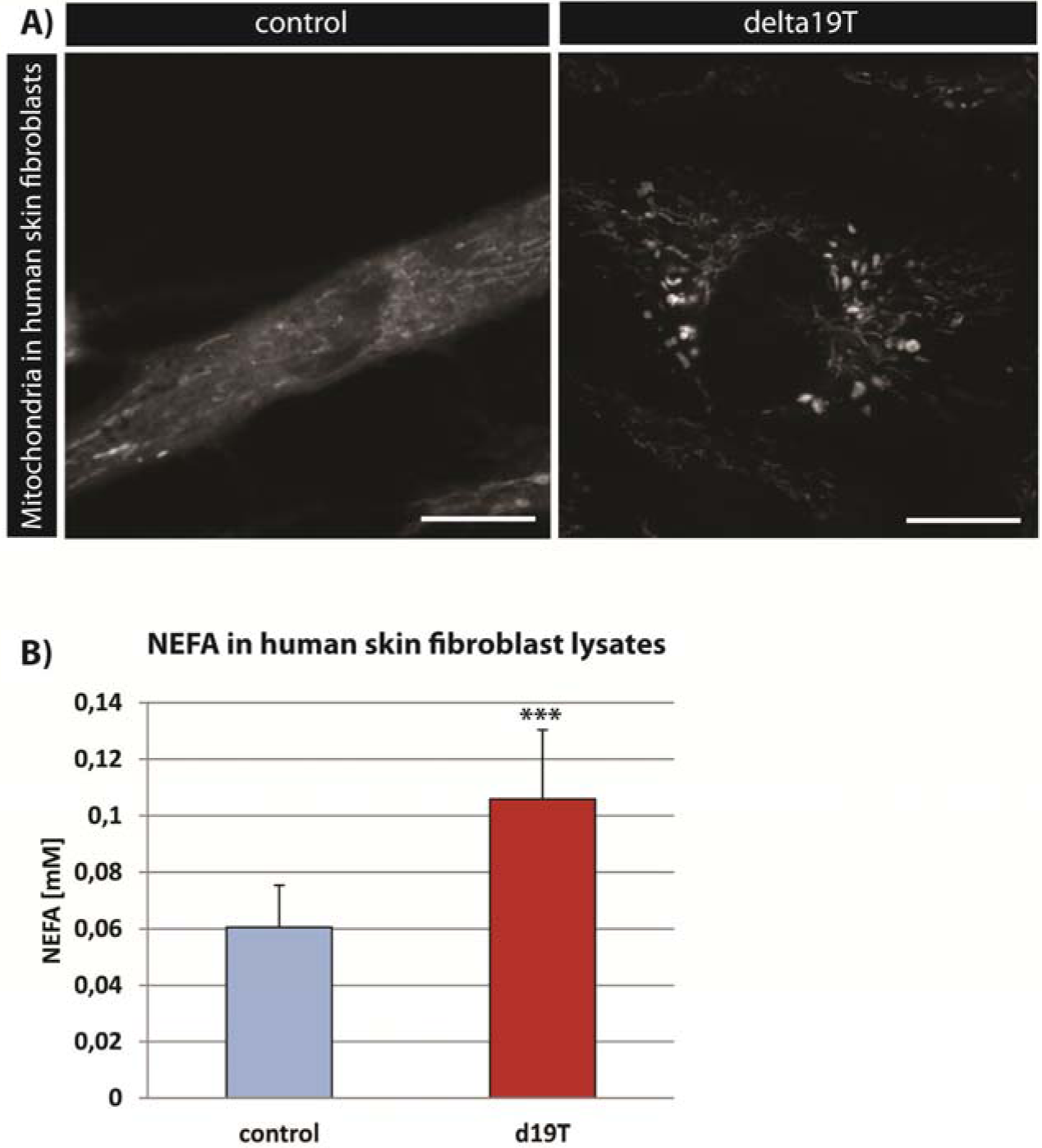
**A)** TMRE staining to visualize active mitochondria in human skin fibroblasts from a healthy person (control) and a patient with a *pex19* mutation (Δ19T). **B)**Amount of non-esterified fatty acids (NEFA) in human skin fibroblasts. Error bars represent SD. ***p<0,001

## Discussion

The tight interconnection of peroxisomes and mitochondria has long been established, and an impact of peroxisome loss on mitochondria has been proposed before. However, the reason for mitochondrial damage as a result of peroxisome deficiency has remained elusive. Here we propose a mechanism by which peroxisome loss due to *pex19* mutation leads to mitochondrial swelling through dysregulated lipolysis. The lipid sensor Hnf4 accounts for upregulation of *lipase 3* and enzymes for mitochondrial acyl-CoA import and β-oxidation. Subsequently, through a vicious cycle of increased lipolysis and NEFA amounts, which in turn can further activate Hnf4 as its ligands, extreme upregulation of *lipase 3*, depletion of the lipid stores, and NEFA accumulation occur, while mitochondrial β-oxidation runs at maximal capacity. This ultimately causes mitochondrial swelling and damage, leading to energy deficiency and lethality. Thus, Hnf4 hyperactivity and dramatic Hnf4-induced *lipase 3* overexpression play major and yet unrecognized roles in the pathophysiology of peroxisome loss.

While *lipase 3* overexpression alone leads to accumulation of toxic NEFA and mitochondrial swelling, which we consider a major contribution to the lethality of the mutant, it does not phenocopy the lethality of *pex19* mutants fully (although it does reduce viability to some extent). This might hint that the peroxisomes, which are still present in *lipase 3* overexpressing animals, but not in *pex19* mutants, play a role in NEFA clearance or mitochondrial turnover rate, thereby counteracting the detrimental effects of dysregulated lipolysis. This hypothesis fits to the fact that peroxisomes and mitochondria share fission and fusion factors and thereby might influence each other’s quality control.

The exact link between Hnf4 and Pex19 remains to be determined. Interestingly, among the putative target genes of Hnf4, several peroxins ( *pex2, pex12* and*pex16*) can be found [26]. This opens up the possibility of a regulatory feedback between peroxisome activity or abundance and Hnf4 activity, which could explain the fact that, in the absence of peroxisomes, Hnf4 activity gets out of hand and initiates a program of maximal lipolysis and beta-oxidation with detrimental results for mitochondria and the whole organism.

## Methods

### Flywork

The *pex19* mutant was generated by imprecise excision following *D. melanogaster* standard techniques. The line *pex19^ΔF7^* was chosen from a jump-out screen and tested as a transcript null. To detect homozygous animals, they were crossed with a CyO-twi-GFP marker. As control flies we used the strain w^1118^(Bloomington stock #3605). Wildtype and heterozygous *pex19^ΔF7^* flies were reared on standard fly food. *hnf4^Δ^33* and *hnf4^Δ17^* flies were kindly provided by Carl Thummel. *hnf^/Δ17^ pex19^ΔF7^* and *hnf4^Δ33^ pex19^ΔF7^* double mutants were generated by genomic recombination, and *hnf4^Δ33/+^ pex19^ΔF7/ΔF7^* mutants by crossing the double mutant with *pex19^ΔF7^*. For survival assays, larvae were collected as 1^st^ instars and transferred to fresh plates with yeast paste. 25 larvae were collected for each condition, and at least 5 independent experiments were conducted. The number of surviving pupae, adults including pharates and viable adults (survivors), which were able to move and lived at least 24h, was counted.

### Cell Culture

Human fibroblast control and Δ19T cells were kept in Dulbecco’s modified eagle medium (DMEM, Gibco) with 10% FBS, 10000 units of Penicillin and 10 mg Streptomycin per ml. For NEFA measurement, 1x10^5^ cells were seeded in 6 well plates and harvested after 48 h. Cells were pelleted and cell pellets were treated like larval tissue (see NEFA section). For stainings, cells were seeded into 8-well-slides for microscopy and stained after 48 h with TMRE and analyzed immediately.

### Imaging

Antibodies used were anti-GFP (Sigma) and anti-Spectrin (DSHB). Secondary antibodies coupled to Alexa dyes were from Molecular probes. Stainings were analyzed using a Zeiss LSM 710 confocal microscope. For ultrastructural analysis, embryos were embedded in 2 % agarose and post fixed with 5 % glutaraldehyde for 2 h at room temperature. Ultra-thin sections were prepared and analyzed using a Zeiss Libra 120 electron microscope. For apoptosis assays, brains of adult flies were dissected and stained using an AnnexinV-FITC apoptosis detection kit (Sigma Aldrich) according to the manufacturer’s instructions.

For immunohistochemistry, we dissected tissue of interest from 3rd instar larvae. Tissue was fixed for 30 minutes in 3,7 % formaldehyde and washed with PBT before and after incubation with primary antibody and Alexa dye-coupled secondary antibody. Tissue was mounted in Fluoromont G and analyzed using a Zeiss LSM 710 confocal microscope. For stainings of neutral lipids with OilRed O, larval tissue was dissected in PBS by inverting the cuticle to provide access to the oenocytes. Tissue was fixed for 20 min in 3,7 % formaldehyde and washed with PBS. Before and after staining with a 60 % OilRed O solution for 30 min, tissue was incubated for 5 min with 60 % isopropanol. Tissue was washed with PBS, mounted in glycerol and immediately analyzed using an Olympus AX70 microscope.

For staining of mitochondria, 96 h old larvae or 5 day old adults were dissected in ice cold PBS, and their Malpighian tubules were stained for 20min at RT with 50nM TMRE (Sigma-Aldrich) in PBS or MitoSOX^TM^ (Molecular Probes) according to the manufacturer’s protocol. The Malpighian tubules were then directly mounted in Fluoromount G and analyzed with a Zeiss LSM 710.

### Lipid profile

For quantification of fatty acid methyl esters (FAME), 15 3rd instar larvae were homogenized in 1N MeHCl in a Precellys 24 homogenizer (peqlab). A minimum of n=7 was analyzed for each condition. C15:0 and C27:0 standards were added and samples were incubated for 45 min at 80 °C. Methylesters were collected by addition of hexane and a 0,9 % NaCl solution. The hexane phase was collected in a new glass vial and concentrated by vaporization. Samples were analyzed by gas chromatography / mass spectrometry using an Agilent HP 6890 with a HP-5MS column.

### Free Fatty Acids

Non-esterified fatty acids (NEFA) were measured by an adaptation of the copper-soap method[38]. In brief, three 3rd instar larvae were weighed and homogenized in 20 µl of 1M phosphate buffer per mg tissue. 25 µl of the supernatant were transferred to 500 µl of Chloroform/Heptane 4:3, and lipids were extracted by shaking the vial for 5 min. Unspecific background provoked by phospholipids was circumvented by addition of 23 mg of activated silicic acid. 300 µl of the chloroform phase were transferred to 250 µl of Cu-TEA (copper-triethanolamine). After shaking and centrifuging, 150 µl of the organic phase were transferred to fresh cups. Liquid was evaporated in a 60 °C heat block, and lipids were dissolved in 100 µl of 100% ethanol. Copper was detected by complexation with a mixture of dicarbazone - dicarbazide, and the colour intensity was measured in a 96 well plate at 550 nm in a TECAN platereader.

### β-oxidation measurements

6 larvae per genotype were washed with PBS and their weight was recorded for normalization purposes. The larvae were inverted in ice cold PBS and permeabilized in ice cold BIOPS buffer (2.77mM CaK_2_EGTA, 7.23mM K_2_EGTA, 5.77mM Na_2_ATP, 6.56mM MgCl_2_.6H_2_O, 20mM taurine, 15mM Na_2_.phosphocreatine, 20mM imidazole, 0.5mM DTT, 50mM MES) containing 100ug/mL saponin (fresh) at 4°C with gentle rocking for 10 mins. Then the larvae were equilibriated in respiration medium (MiR05, 0.5mM EGTA, 3mM MgCl2.6H2O, 60mM K-Lactobionate (lactobionic acid is dissolved in H2O and pH is adjusted to pH 7.4 with KOH), 20mM Taurine, 10mM KH2PO4, 20mM HEPES, 110mM sucrose, 1g/L fatty acid free BSA) supplemented with 0.5mM carnitine. The larvae were added into the oxygraph chambers and oxygen concentration was brought to around 500uM by using catalase and H_2_O_2_. After basal respiration was recorded, 5uM palmitoylCoA was added to the chamber. Fatty acid β-oxidation was induced by adding complex I substrates, ETF (electron transfer flavoprotein) substrates, and ADP (10mM proline, 10mM pyruvate, 5mM malate, 5mM glutamate, 2mM ADP and 15mM glycerol-3- phosphate). After that, etomoxir was added at the indicated concentrations to block fatty acid transfer into mitochondria via CPT1, thereby blocking β-oxidation and leaving complex I–dependent respiration. Finally, residual oxygen consumption (ROX) was measured by inhibiting complex III with antimycin A. All values were corrected for ROX. β-oxidation was calculated by subtracting etomoxir-resistant respiration from respiration in the presence of all substrates.

### Realtime qPCR

Whole RNA of 3rd instar larvae was isolated using TriFast reagent (peqlab). Tissue was homogenized using a Precellys 24 homogenizer (peqlab). Transcription to cDNA was performed using the Quantitect Reverse Transcription Kit (Quiagen). Quantitative PCR was performed with a CFX Connect cycler (biorad). Each experiment was repeated at least 5 times.

## Author contribution

Conceptualization, M.H.B., J.S., C.W. and M.H.; Methodology, M.H.B., J.S., C.W., D.G.; Investigation, M.H.B., J.S., C.W., D.G., D.S.; Writing – Original Draft, M.H.B. and J.S.; Writing – Review & Editing, M.H.B., J.S., M.H. and A.A.T.; Visualization: M.H.B. and J.S.; Funding Acquisition, M.H. and A.A.T., Supervision, M.H. and A.A.T.

## Acknowledgments

We thank the Bloomington stock center for fly strains, Carl Thummel for providing Hnf4 fly lines and Ingo Zinke and Michael Pankratz for the ppl-Gal4 and UAS lip3 line. We thank Christoph Thiele for input and help with the NEFA measurements. We thank Gabriele Dodt, Ronald Wanders, Heike Schulze and Konrad Sandhoff for human fibroblast lines, Mélisande Richard for help with the ultrastructural analysis, Fatmire Bujupi for help with the characterization of the *pex19* mutant, the group of Peter Dörmann for help with mass spectrometry analysis of FAMEs, and members of the Hoch group for discussion.

The work was funded by grants to M.H. *(SFB 645, TPB1 and TR 83, TP A7)* and by the Helmholtz Portfolio grant “*metabolic dysfunction*” to M.H. and A.A.T. M.H. is a member of the Bonn Excellence Cluster *ImmunoSensation.*

